# The functionally plastic rod photoreceptors in the simplex retina of Little skate (*Leucoraja erinacea*) exhibit a hybrid rod-cone morphology and enhanced synaptic connectivity

**DOI:** 10.1101/2023.06.28.546621

**Authors:** Laura Magaña-Hernández, Abhiniti S. Wagh, Jessamyn G. Fathi, Julio E. Robles, Beatriz Rubio, Yaqoub Yusuf, Erin E. Rose, Daniel E. Brown, Priscilla E. Perry, Elizabeth Hamada, Ivan A. Anastassov

## Abstract

The retinas of the vast majority of vertebrate species are termed “duplex” – that is, they contain both rod and cone photoreceptor neurons in different ratios. The retina of Little skate (*Leucoraja erinacea*) is a rarity among vertebrates because it contains only rod photoreceptors and is thus “simplex”. This unique retina provides us with an important comparative model and an exciting opportunity to study vertebrate rod circuitry within the context of a functional, evolutionarily optimized system, all without the concern about artifacts from genetically modified rod-only mouse models. Perhaps even more importantly, the Leucoraja retina is able to function under both scotopic and photopic ranges of illumination with a single complement of photoreceptors. It is currently unknown what structural characteristics mediate this remarkable functional plasticity. To address this question, we performed serial block-face electron microscopy imaging and examined the structure of rods and their post-synaptic partners. We find that skate rods exhibit ultrastructural characteristics that are either common to rods or cones in other vertebrates (e.g., outer segment architecture, synaptic ribbon number, terminal extensions), or are uniquely in-between those of a typical vertebrate rod or cone (e.g., number of invaginating contacts, clustering of multiple ribbons over a single synaptic invagination). We therefore hypothesize that the unique hybrid rod-cone structure of skate rods and their post-synaptic partners is correlated with the ability of the skate visual system to function across scotopic and photopic ranges of illumination. These findings have the potential to reveal as yet undescribed principles of vertebrate retinal design.

**Significance statement:** The vast majority of vertebrate retinas are duplex and have mixed rod-cone populations of photoreceptors in varying ratios. The processing of visual information in a duplex retina tends to be separated between rod and cone systems, which mediate function under scotopic and photopic lighting conditions, respectively. However, the cartilaginous fish Little skate (*Leucoraja erinacea*) has a simplex retina, comprised solely of rod photoreceptors. Skate rods are also unusual because they have the ability to retain function over a full range of lighting conditions. We have little knowledge about the ultrastructural anatomy of the skate retina, and we hypothesize that this functional plasticity can be traced back to morphological adaptations at the level of individual photoreceptors and the downstream retinal circuitry, thus illuminating new pathways for the processing of visual information among vertebrates.

## Introduction

The Little skate (*Leucoraja erinacea)* is a member of the elasmobranchii subclass of cartilaginous fishes and has been a consequential model system for studies of fin and limb development^1, 2^, skeleton formation ^3^, electroreception ^4^ and even the evolution of walking on land^5^. The skate visual system has received a lot less attention, but appears to be no less fascinating. For example, the neural retina of this animal goes against the trend of the vast majority of other vertebrate retinas and appears to be comprised of only rod photoreceptors ^6, 7^. A mixed rod-cone photoreceptor retina is a lot more typical among vertebrates, where rods and cones are found in different ratios^8^, and where rods mediate scotopic (dim light) vision, while cones mediate photopic (daylight) vision^9^. As such, the pure-rod nature of the skate retina has provided a unique opportunity to study a naturally occurring cone knock out visual system^10^, unencumbered by some of the artifacts of genetic manipulation in rodents^11, 12^. Perhaps even more surprising, however, is the fact that the skate retina can function under both scotopic and photopic light conditions with a monotypically pure photoreceptor system^13^. A number of elegant classical studies by Dowling and Ripps have shown that under scotopic conditions, skate rods can function at the theoretical threshold of sensitivity and detect single photons^13–15^. But, after a relatively brief period of light-adaptation, they speed up their kinetics, lower their sensitivity, and expand their functional capabilities to photopic levels of illumination^13, 16^. Furthermore, the downstream components of the skate retinal circuitry can also adapt from scotopic to photopic conditions (and back) and continue to transmit the visual message^16, 17^. How this functional plasticity in skate rods and the downstream circuitry happens is still not entirely understood and it is surprising that we also know very little about the ultrastructural anatomy and connectivity of skate retinal neurons.

To the best of our knowledge, only two studies have examined any aspects of the ultrastructure of neurons in the skate retina in any appreciable detail. Szamier and Ripps (1983)^18^ used conventional electron microscopy to examine the juvenile skate retina and describe the disk shedding properties of skate rods, showing that what appeared to be cone-like cells in younger animals were in fact immature rods. In a separate study, Malchow and colleagues^19^ (1990) examined the ultrastructural and functional properties of two types of skate horizontal cells and showed that they are physiologically and anatomically distinct. A somewhat surprising finding, given the all-rod nature of the skate retina and the tendency of mixed rod-cone retinas to dedicate more horizontal cell types to cone processing, rather that rod processing^20^.

Recent advances in 3D ultrastructural imaging have provided us with an opportunity to resolve and examine fine details of the skate retina anatomy and to begin assembling a connectome of this unique visual system. In the present study, we used serial block-face scanning electron microscopy method (SB-3DEM), a cutting-edge method for anatomical reconstruction, to examine the ultrastructure of skate rods and the post-synaptic processes invaginating into rod terminals. We show that skate rods display structural elements that are commonly found in other vertebrate rods, mixed with elements that are either more typical of vertebrate cones, or completely unique to skate rods. We suggest that skate rods possess a hybrid rod-cone architecture, which likely mediates the ability of the skate visual system to function across scotopic and photopic ranges of illumination. These findings have the potential to significantly expand our understanding of the vertebrate visual system and reveal as yet undescribed principles of vertebrate retinal design.

## Methods

### Animals

All animal procedures were approved by the respective Animal Care and Use Committees at the Marine Biological Laboratory (Woods Hole, MA) and San Francisco State University (San Francisco, CA). Wild caught adult and in-house bred juvenile Little and Winter skate animals (*Leucoraja erinacea* & *Leucoraja ocellata*) were obtained from the Marine Biological Laboratory Marine Resources Center (Woods Hole, MA). They were kept in a recirculating seawater system at 12-13°C. Circulating seawater was subjected to continuous physical, biological and chemical filtration. Animals were kept under a 12/12 hr. light/dark cycle and fed finely chopped frozen squid and mysids once a day. Both species (*L. erinacea* & *L. ocellata)* were used in this study, as they are closely related, frequently co-habit in the wild, and extensive studies have shown no anatomical or physiological differences between the retinas of either species^6, 7^. Animals were monitored daily for health and signs of general distress and all studies were performed following animal euthanasia. Prior to euthanasia, animals were anesthetized with sodium bicarbonate-buffered 0.02% tricaine methanesulfonate (Syndel, Canada) until unresponsive, followed by fast cervical transection and pithing. This method of euthanasia is consistent with the American Veterinary Medical Association (AVMA) Guidelines on Euthanasia.

### Tissue preparation

Retinal tissue was harvested following euthanasia from the eyes of 3 adult skates. Eyes were enucleated under ambient illumination; the cornea and lens were removed and the vitreous drained. The retina was left attached to the choroid and cartilaginous sclera to protect it from structural damage and aid in subsequent sectioning. The resulting eyecups were immediately fixed with 4% Paraformaldehyde + 2.5% Glutaraldehyde in 0.1 M Cacodylate buffer (pH 7.2) for 5-7 days at 4°C. Fixed samples were shipped on ice to the 3DEM Ultrastructural Imaging and Computation at the Cleveland Clinic Lerner Research Institute (Cleveland, OH), where tissue were subjected to post-fixation with OsO_4_, graded dehydration with ethanol, *en-bloc* staining with uranyl acetate, and infiltration and embedding in epoxy resin.

### Imaging

Imaging of samples and collection of raw data were performed off-site at the Cleveland Clinic 3DEM Ultrastructural Imaging and Computation Core (Cleveland, OH). Large volume 3D electron microscopy was performed on retinal pieces embedded in epoxy resin using the serial block-face scanning electron microscopy method (SB-3DEM). A Teneo Volumescope system (Thermo Fisher Scientific, Waltham, MA) and a Zeiss Sigma VP system (Carl Zeiss Microscopy GmbH, Jena, Germany) equipped with a Gatan 3View in-chamber ultramicrotome stage, were used to image the different samples. Samples were sectioned and imaged in the cross-sectional orientation, which allowed for the visualization of all retinal layers and cell types. The datasets used and analyzed in the present study are from a region of interest in the outer plexiform layer (P2R9 volume) and from a full cross-section of the retina (HVMS volume). The region of interest dataset had a width and height of 27.6μm and a depth of 21.5μm; voxel size was 4.5×4.5×70nm (xyz). The full cross-section dataset had a width of 88μm, a height of 304μm and a depth of 22μm; section thickness is 0.075μm; voxel size was 10×10×75nm (xyz).

### Data analysis and statistical procedures

Segmentation, 3D reconstructions, surface area and volume measurements were obtained with Reconstruct software^21^. All other quantitate measurements were obtained with Amira software (Thermo Fisher Scientific, Waltham, MA). Quantitative and statistical analyses were completed with Prism software (GraphPad Software, La Jola, CA). Two-tailed unpaired t-tests with and without Welch’s correction were used for comparisons of rod OS and IS area, volume and length. OS and IS diameter comparisons parameters were done with a two-tailed Mann-Whitney test. P-values and replicates are listed in the figure legends and main text.

## Results

### The simplex retina of Leucoraja erinacea contains only rod photoreceptors

The elasmobranch fish *Leucoraja erinacea* (common name, Little skate) is a benthic species commonly found off the US East Coast. A juvenile hatchling animal is about 80-100mm long, tip to tail, with a disc diameter of 4-5mm (Fig. 1A). A very closely related species, *Leucoraja ocellata* (common name Winter skate), naturally co-habits in the same waters as *L. erinacea* and is morphologically identical. The pupil of the skate eye is covered by a structure called the *operculum pupillare*^22, 23^. The *operculum pupillare* completely covers the pupil when the animal is light-adapted (Fig. 1B), and completely retracts to expose the whole pupil when the animal is dark-adapted (Fig. 1C). Both species have retinas that are termed “simplex” and contain only rod photoreceptors (Fig. 1D). Numerous studies in both species have confirmed that their retinas are identical and pure-rod^6, 7, 24^. Furthermore, skate rods exhibit a kind of “functional plasticity”, which allows them to seamlessly adapt to both scotopic and photopic illumination conditions^13, 14, 25^. This functional trait appears to be conserved and neurons that are downstream of rods can also adapt from scotopic to photopic conditions (and back) and continue to transmit the visual message^14, 16, 26^. For the purposes of this study, and for the reasons stated earlier, we have not differentiated between the two species.

**Figure 1.**
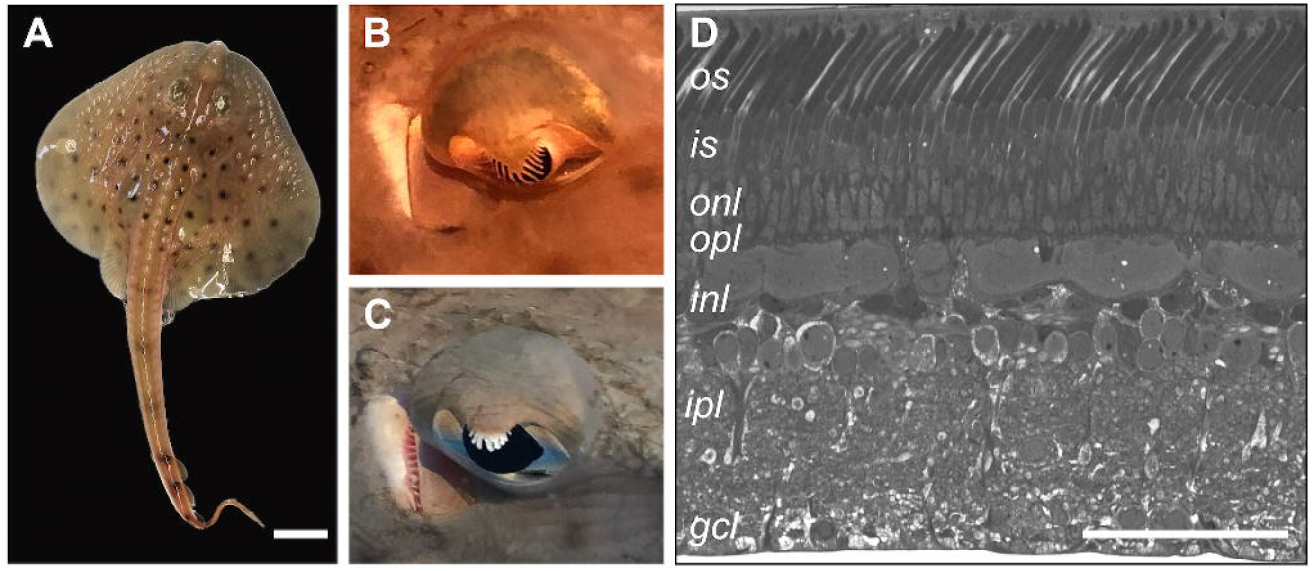
The retina of the Little skate is simplex and only rod photoreceptors can be distinguished by gross morphology. **A)** A juvenile hatchling example of the Little skate (*Leucoraja erinacea*). The skate is a benthic fish found off the US East Coast. It co-habits naturally with another species, *Leucoraja ocellata* (Winter Skate) and both have virtually identical appearance. Their retinas are also both simplex and pure-rod (scale bar = 1cm). **B)** The iris operculum of a light-adapted skate eye can be seen covering the majority of the cornea. **C)** During dark-adaptation, the iris operculum completely retracts and allows for a maximum amount of light to enter the eye. The spiracle, a small, round opening, is immediately posterior to the eye (to the left of the eye in both B & C). Spiracles are used for respiration and draw water into the gill chambers. The gill slits are located on the ventral side of the animal. **D)** A histological cross-section of the adult skate retina stained with Methylene Blue and Azure II. Only rod outer segments are visible in cross-section, as described previously by Dowling and Ripps (1990). The retina of the Winter Skate is also pure-rod and not distinguishable from that of the Little skate in any appreciable way (scale bar = 100µm). OS - outer segments; IS - inner segments; ONL - outer nuclear layer; OPL - outer plexiform layer; INL - inner nuclear layer; IPL - inner plexiform layer; GCL - ganglion cell layer.

### Serial EM imaging of the L. erinacea retina confirms exclusive presence of rods

We performed serial block-face 3D electron microscopy imaging on several retinal samples from adult skates. To our knowledge, this is the first time that this imaging approach has been applied systematically to any simplex retina. The results presented here were obtained from 2 different datasets: a full cross-section dataset (HVMS) and a high-resolution region of interest (ROI) dataset (P2R9). Figures 2A and 2F show representative 2D images of each dataset. The HVMS dataset is from a full cross-section of an adult retina with a width of 88μm, a height of 304μm and a depth of 22μm. Section thickness is 0.075μm and the voxel size is 10×10×75nm; a virtual stack of the volume with dimensions can be seen in Fig. 2B and 2D. The P2R9 dataset is from an ROI in the outer plexiform layer (OPL) and has a width and height of 27.6μm and depth of 21.5μm; voxel size is 4.5×4.5×70nm (Fig 2G and 2I). Manual and semi-manual segmentation in the outer retina from the HVMS dataset allowed us to obtain 3D reconstructions of whole rods, including inner and outer segments (IS and OS), synaptic terminals, and a significant portion of invaginating post-synaptic processes (Fig. 2C1 and 2C2). The full volume of the HVMS dataset (with raw data excluded) and the reconstructions of 9 full rods, along with some of their connecting postsynaptic processes, can be seen in Fig. 2D and Fig. 2E1-E3. Skate rods take up about 50% of the cross-sectional length of the whole retina. Reconstructing 9 rods and a portion of their connected post-synaptic dendritic architecture covered the full z-dimension of the HVMS dataset suggesting that a depth of 22μm is sufficient for partial reconstruction of post-synaptic architecture, but likely insufficient for the reconstruction of full postsynaptic partners, like entire bipolar or horizontal cells, which appear to be quite a lot larger than the HVMS dataset spans in the *z*-plane. Nevertheless, we believe we have sufficient data to differentiate between different postsynaptic processes and assign them to putatively different post-synaptic rod partners. This estimation is confirmed by the significantly different morphology and spatial location of rod post-synaptic processes reconstructed from the high resolution P2R9 dataset (see Fig. 2J1, 2J3 and 2J4). Fine details of rod synaptic architecture, like ribbons synaptic vesicles and invaginating contacts can be distinguished readily from the high-resolution ROI dataset P2R9, a representative 2D image of which is shown in Fig. 2F. Segmentations from different neighboring rod terminals and the post-synaptic processes invaginating into each terminal can be seen in Fig. 2H1 and 2H2. Note the marked and readily distinguishable synaptic ribbons and ribbon-docked synaptic vesicles in Fig. 2H2 and 2J2. The full R2P9 volume allowed for a full reconstruction of 20 rod terminals and partial reconstruction of another 9 rod terminals (Fig. 2I and 2J2). Synaptic ribbon clusters could be used to determine the location of an individual rod terminal (Fig. 2J1 and Fig. 4C3).

**Figure 2.**
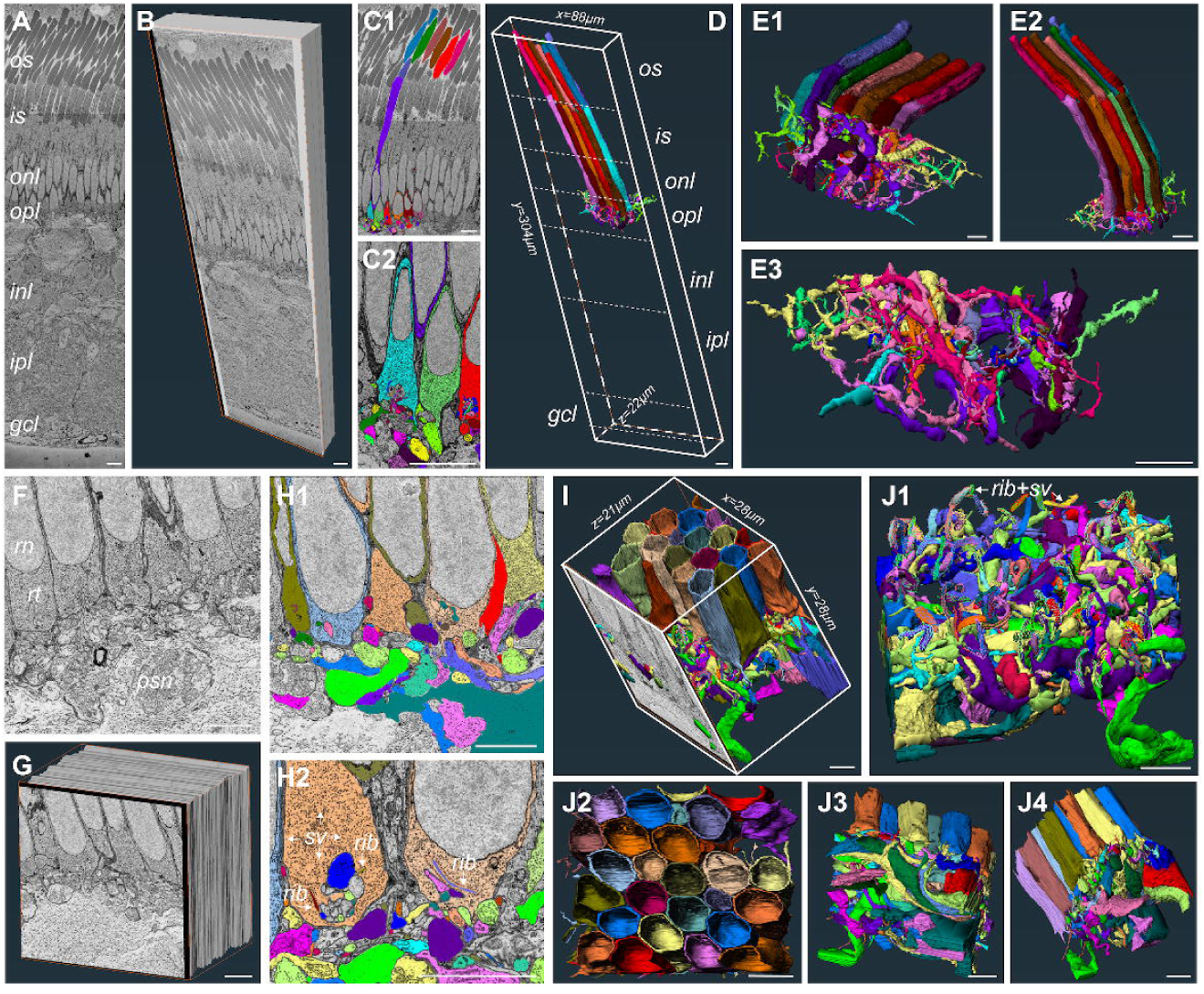
Serial block-face scanning electron microscopy (SB-3DEM) imaging and segmentation of skate retinal tissue. A) Single section from full cross-section HVMS dataset. B) A virtual block reconstruction of all sections within volume (scale bars = 10µm). C1-C2) Examples of semi-manual segmentation of structures using Reconstruct software (scale bars = 10µm). D) 3D reconstructions of rod photoreceptors and where they fall within the EM volume, and the dimensions of the HVMS dataset (scale bar = 10µm). E1-E3) Side and bottom-up views of 3D reconstructions of entire rods and their associated post-synaptic dendritic processes (scale bars = 10µm). F) Single section from full cross-section P2R9 dataset, rt - rod terminal; rn - rod nucleus; psn - post-synaptic neuron, rib – synaptic ribbon, sv – synaptic vesicles (scale bar = 5µm) G) A virtual block reconstruction of all sections within volume (scale bar = 5µm). H1-H2) Examples of semi-manual segmentation of structures within P2R9 volume using Reconstruct software (scale bars = 5µm). I) 3D reconstructions of rod photoreceptor terminals and their associated post-synaptic dendritic processes and the dimensions of the volume; one section of the raw data is also shown. J1) 3D reconstructions of post-synaptic processes with rod terminals removed and rod synaptic ribbons with docked vesicles displayed (J2-J4) Side and top-down views of all 3D reconstructions within volume (scale bars = 5µm).

### The outer segments of skate rods display separated stacked membrane morphology, which is typical for rods of duplex retinas

The skate rod cell is long, which appears quite similar to the anatomy of mammalian duplex retina rods^27, 28^, but not quite as slender and with a larger OS/IS diameter, which is more typical of the anatomy of rods in non-mammalian duplex retinas^29, 30^. Rods take up ∼50% of the cross-sectional length of the retina and there are clearly distinguishable outer segments (OS), inner segments (IS) and synaptic terminals (ST) (Fig. 3A1 - 3A4). Additional rod features are described in the sections that follow. High-resolution single image TEM data from Szamier and Ripps (1983)^18^ has shown previously that the OS of *L. erinacea* rods have the typical ordered stacks of internal membrane disks (in all likelihood holding the proteins that take part in the rhodopsin light response cascade), which are physically separated from the rod outer membrane. Although not of the same high resolution as conventional 2D TEM, our SB-3DEM data allowed us to readily confirm the finding of Szamier and Ripps of stacked internal disks separated from the rod outer membrane (Fig. 3B1 and 3B2). There is a slight offset in the location vertical location of each OS, which can be seen in Fig. 3B3 and 3B4.

**Figure 3.**
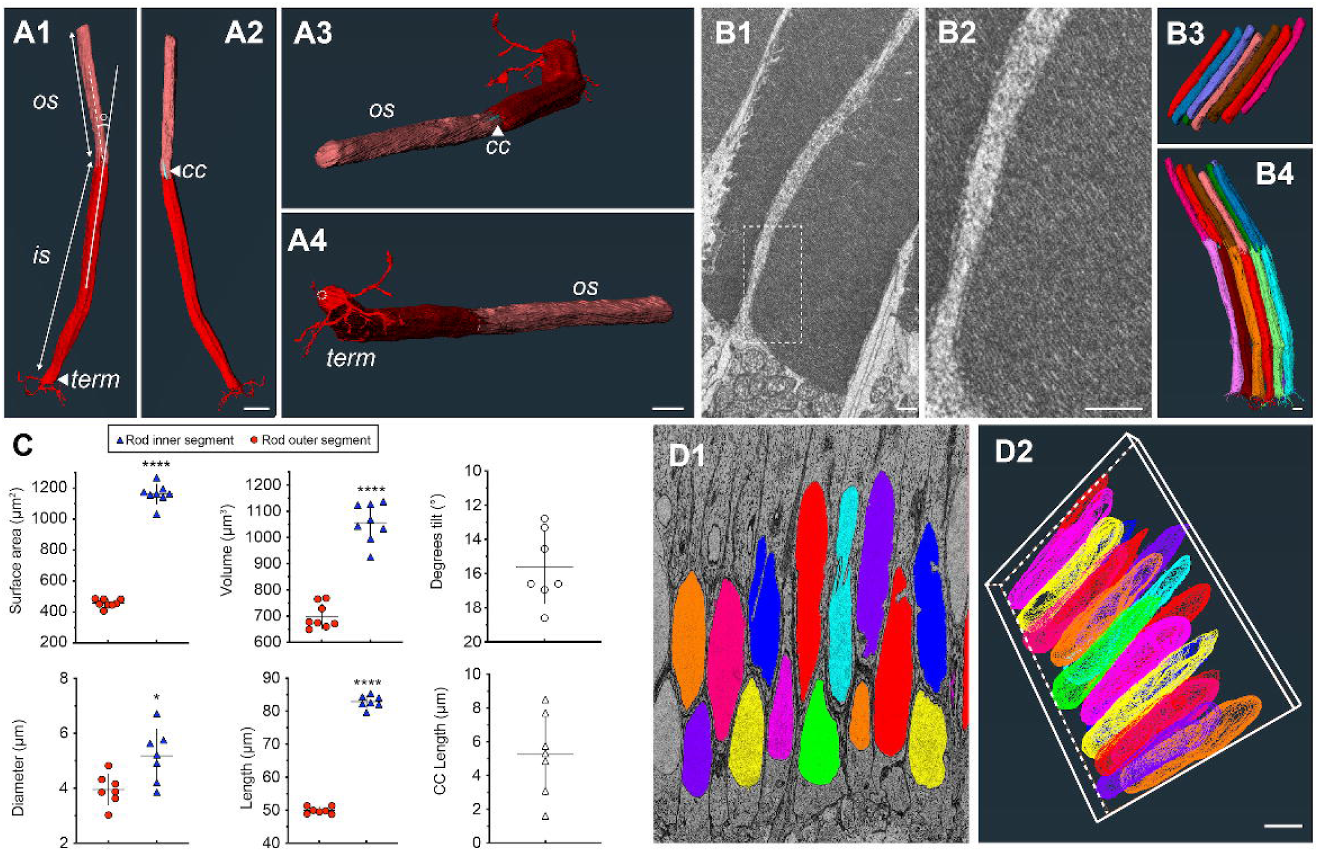
Anatomy and ultrastructure of whole skate rods. A1-A4) Reconstructions of a single rod showing outer and inner segment (OS, IS), tilt angle between OS and IS, connecting cilium (CC) and synaptic terminal (term) with telodendria. The opening through which invaginating processes enter the rod terminal is off-set from the center and is indicated with a dashed circle in A4 (scale bars =10µm). B1-B2) Disc membrane stacking and separation of membrane discs from outer cell membrane is readily seen in the skate rod OS, which is typical of vertebrate rods (scale bars = 1µm). B3-B4) Relatively little stacking of outer segments or whole rod cells is seen, as opposed to murine or primate retinas (scale bars = 1µm). C) Quantifications and comparisons of rod OS and IS surface area (unpaired t-test, two-tailed, p<0.0001, n=8), diameter (unpaired Mann-Whitney test, two-tailed, p=0.0186, n=7), volume (unpaired t-test, two-tailed, p<0.0001, n=8), length (unpaired t-test, two-tailed, p<0.0001, n=7), OS vs. IS tilt angle, and CC lengths of fully reconstructed cells. D1-D2) Relatively little stacking of rod nuclei is seen, when compared to mammalian (e.g. mouse or rabbit) duplex retinas (scale bars = 5µm).

### Rod outer and inner segments vary little in diameter from each other, inner segment is consistently longer and larger, and nuclei display moderate stacking

We performed detailed quantitative analysis of different rod features based on the 3D reconstructions obtained. These analyses showed that inner segments were consistently longer than outer segments by ∼40% (mean of 83μm IS vs. 50μm OS, Fig. 3C). Consistent with this observation, surface area (mean of 1162μm^2^) and volume (mean of 1056μm^3^) for IS were also significantly larger than for OS (457μm^2^ and 699μm^3^, respectively, Fig. 3C). Diameter of IS (measured as an average of diameters at 3 different points along the segments) was only moderately, although still significantly, bigger (Fig. 3C). We also measured the tilt angle between the IS and OS of each reconstructed rod and each cell showed a consistent mean tilt angle of 15.6° (see angle measurement example in Fig. 3A1 and tilt angle values in Fig. 3C). This falls well within the values of tilt angle measurements for skate rods within the visual streak, obtained recently by Mäthger and colleagues^31^. Tilt angle values are also in agreement with the fact that tissue for SB-3DEM imaging was taken from the tapetal area of the retina, which often overlaps with the visual streak in elasmobranchs^22^. Interestingly, there is little stacking of photoreceptor nuclei in the skate retina, unlike what is often observed in a number of mammalian species^32–34^, where rod nuclei are stacked in columns of up to 12 or 14. However, reconstructions from a number of skate rod nuclei reveal that only moderate stacking is present (Fig. 3D1 and 3D2). This appears to be a fundamentally different architecture from that of rods in mammalian retinas of commonly used murine model organisms^35, 36^.

### Rods have multiple ribbons, which are centered in clusters around a single terminal invagination

We continued our investigation of rod morphology in the skate by focusing on a common feature in the terminals of primary sensory neurons: synaptic ribbons. These organelles serve as organizing centers that tether synaptic vesicles at the active zones of photoreceptor and bipolar cell terminals in the vertebrate retina ^37, 38^. Synaptic ribbons in the retina have been studied extensively ^39–41^ and in mammals the synaptic terminal of vertebrate rod spherules tends to contain one ribbon centered around a single terminal invagination^42–44^. In some fishes and amphibians, however, rods can contain more than one ribbon^30, 45^. Mammalian cone pedicles, on the other hand, tend to contain multiple ribbons and invaginations, where each ribbon is centered over its dedicated invagination, but all are contained in the same terminal^46, 47^. Teleost and amphibian cone pedicles vary and can sometimes have a single large invagination surrounded by multiple ribbons^48^. Surprisingly, skate rod terminals exhibit a morphology that is in-between that of a typical vertebrate rod and cone. That is, there is a single invagination with multiple ribbons centered around it (Fig. 4A1-A2). Furthermore, the number of ribbons clustered over the invagination is not constant and we repeatedly encountered terminals with either 1, 2, 3 or 4 ribbons (Fig. 4B1-B4). Ribbon clusters assume a spherical arrangement over the invaginating post-synaptic processes, covering it like an umbrella (Fig. 4C1-C2). We examined 74 rod terminals across the HVMS and P2R9 datasets (as well as a third partial ROI dataset not shown here) and found that ribbon distribution is heavily skewed towards terminals with two (n=32) or three ribbons (n=36), with one or four ribbons (n=3, for both) at the tail end of the distribution and a relatively rare occurrence (Fig. 4D). The spatial arrangement of terminals with different number of ribbons did not show any appreciable pattern (Fig. 4C3-C4) and a correlation between the number of ribbons and the number if invaginating processes is largely lacking (see Fig. 7G).

**Figure 4.**
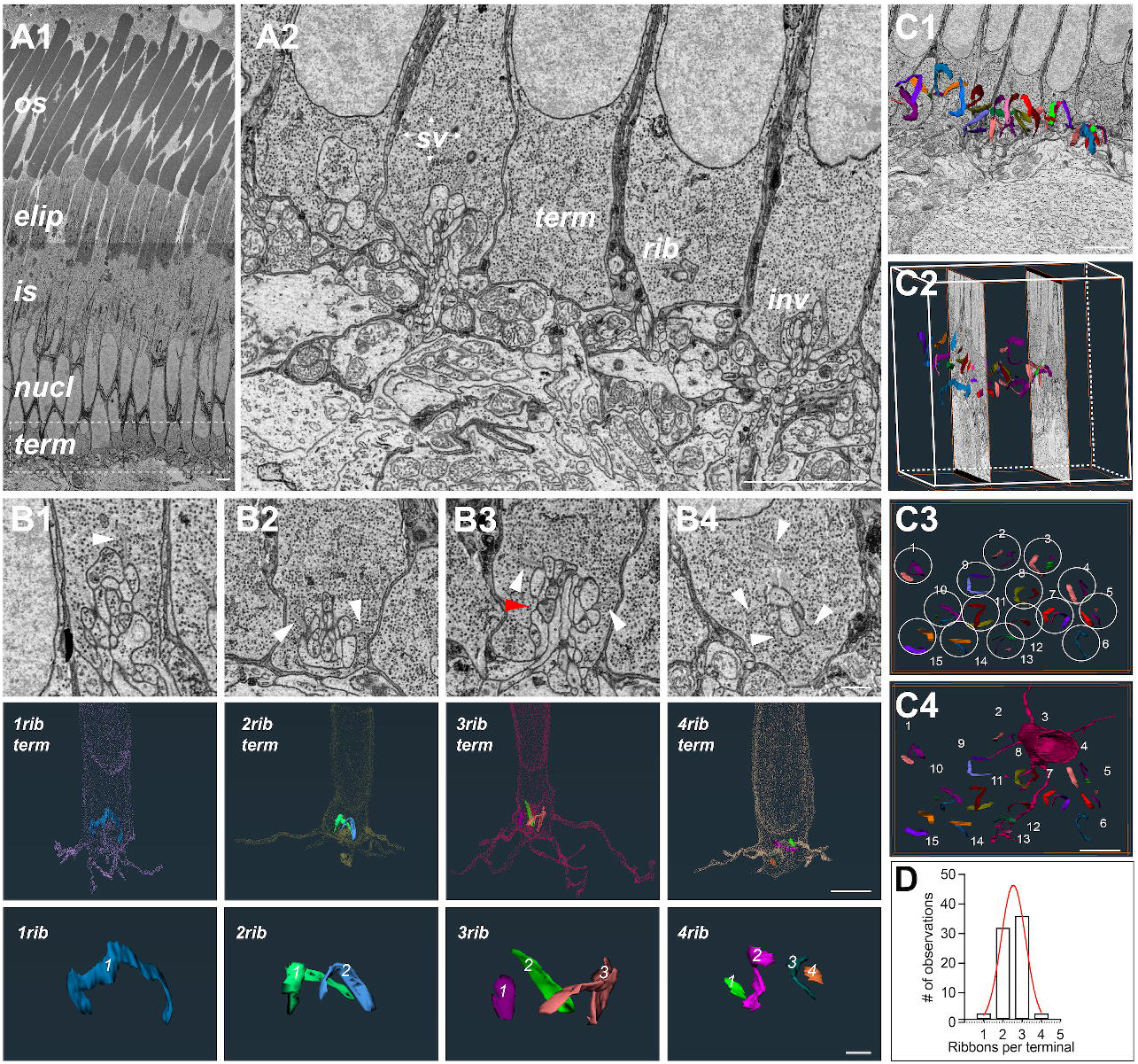
Skate rod terminals have multiple synaptic ribbons. A1) A single EM image from HVMS dataset of the outer skate retina with the different areas of photoreceptors indicated; os – outer segment, elip – ellipsoid (location of mitochondrial clustering), is – inner segment, nucl – nucleus, term – synaptic terminal (scale bar = 5 µm). A2) Close-up of 5 terminals (from left to right) from the P2R9 dataset showing some of the anatomy typical features found in skate rod terminals, as wells as some of the post-synaptic processes that invaginate into those terminals; sv – synaptic vesicles, term – terminal, rib – synaptic ribbon, inv – invagination (scale bar = 5 µm). B1) An EM image with an example (white arrowhead) of a single ribbon terminal (top); a 3D reconstruction of the single ribbon (bottom). B2) An EM image with an example (white arrowheads) of a two ribbon terminal (top); 3D reconstruction of the same terminal with the two ribbons visible inside it (middle); 3D reconstruction of both ribbons together (note the appearance of a spherical arrangement of the ribbons over the single invagination). B3) An EM image with an example (white arrowheads) of a three ribbon terminal. The red arrowhead shows the approximate location of the third ribbon, which appears in later sections (top); 3D reconstruction of the same terminal with the three ribbons visible inside it (middle); 3D reconstruction of all three ribbons together (note again the appearance of a spherical arrangement of the ribbons over the single invagination. B4) An EM image with an example (white arrowheads) of a four ribbon terminal (top); 3D reconstruction of the same terminal with the four ribbons visible inside it (middle); 3D reconstruction of all four ribbons together. Scale bars in EM images = 1µm; in terminal reconstruction images = 5µm, and in ribbon images = 1µm. C1-C2) Ribbons appear in clusters and can be seen here overlaid with individual raw EM images in 2D (C1) and 3D (C2). C3-C4) Rod ribbons assume a “spherical” arrangement - ribbons belonging to an individual rod terminal are circled and numbered. An example terminal belonging to the ribbons in circle 3 is shown. Note widely extending telodendria (scale bars = 5µm. F) Rod terminals have between 1 and 4 ribbons, which is quantified here in the histogram of ribbon distributions in the data to date.

### Rod terminals have multiple, cone-like telodendria that extend to form a meshwork

The terminals of vertebrate rod and cone photoreceptors, in particular mammalian ones, differ from each other in a number of morphological features, one of them being the telodendria extending from the cone pedicles^49^. Usually, cone pedicles have multiple long extensions (i.e., telodendria), which may connect them to neighboring cones or other post-synaptic cells within the local circuitry^50^. Rod spherules tend to lack telodendria and connect to each other or other cones via gap junctions^51, 52^. Non-mammalian rods and cones are again more diverse, but it appears that rod telodendria are significantly less numerous and shorter than cone telodendria^53, 54^. However, the unusual terminal morphology of skate rods continued to manifest itself in this regard as well. All rod terminals that we examined and reconstructed for telodendria across our datasets (n=41), had a number of long and extending telodendria, as can be seen from the example segmentation from the raw data in Fig. 5A. 3D reconstructions of telodendria show that they form an intricate meshwork of processes between rods (Fig. 5B1-B2). We analyzed 41 full rod terminals and the distribution of the number of telodendria per rod terminal is shown in Fig. 5D. We encountered terminals with 6 or 7 telodendria most often, while the length of each process - within and between terminals - varied considerably and was not clearly correlated with how many telodendria per terminal were present (see Fig. 6). Putative synaptic vesicles were identified in multiple locations along the length of different telodendria, suggesting that there are multiple synaptic contacts that telodendria make to neighboring processes (Fig. 5C1-C3). These processes appeared to be either other telodendria, or the putative dendrites of post-synaptic cells. A detailed mapping of input and output telodendria contacts is part of a separate study.

**Figure 5.**
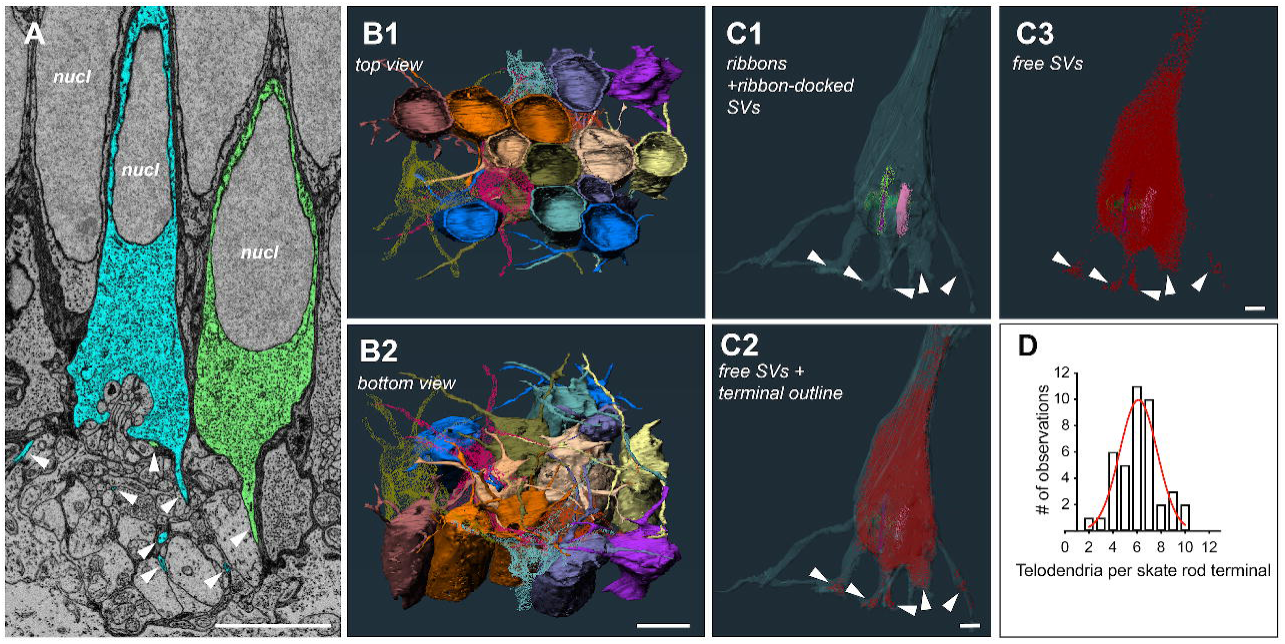
Skate rod terminals have a hybrid terminal morphology, with multiple extending telodendria, similar to mammalian cones. A) Single section from HVMS volume showing segmentation of two neighboring rod terminals and their extending telodendria in cyan and green (white arrowheads; scale bar = 5µm). B1-B2) Top and bottom view of partial rods with reconstructed terminals and the extending telodendria of each. Several of the terminals and their telodendria have been made transparent and not all terminals are shown for clarity (scale bar = 5µm). C1) A transparent 3D reconstruction of a single terminal with ribbons and ribbon-docked vesicles visible, the endings of telodendria are indicated by white arrowheads. C2) All free vesicles within the same terminal. C3) Vesicles are often observed at the ends of telodendria (white arrowheads), suggesting synapses onto other rods or second-order cells (scale bar = 1µm). D) Distribution of telodendria numbers from all reconstructed terminals across both datasets.

**Figure 6.**
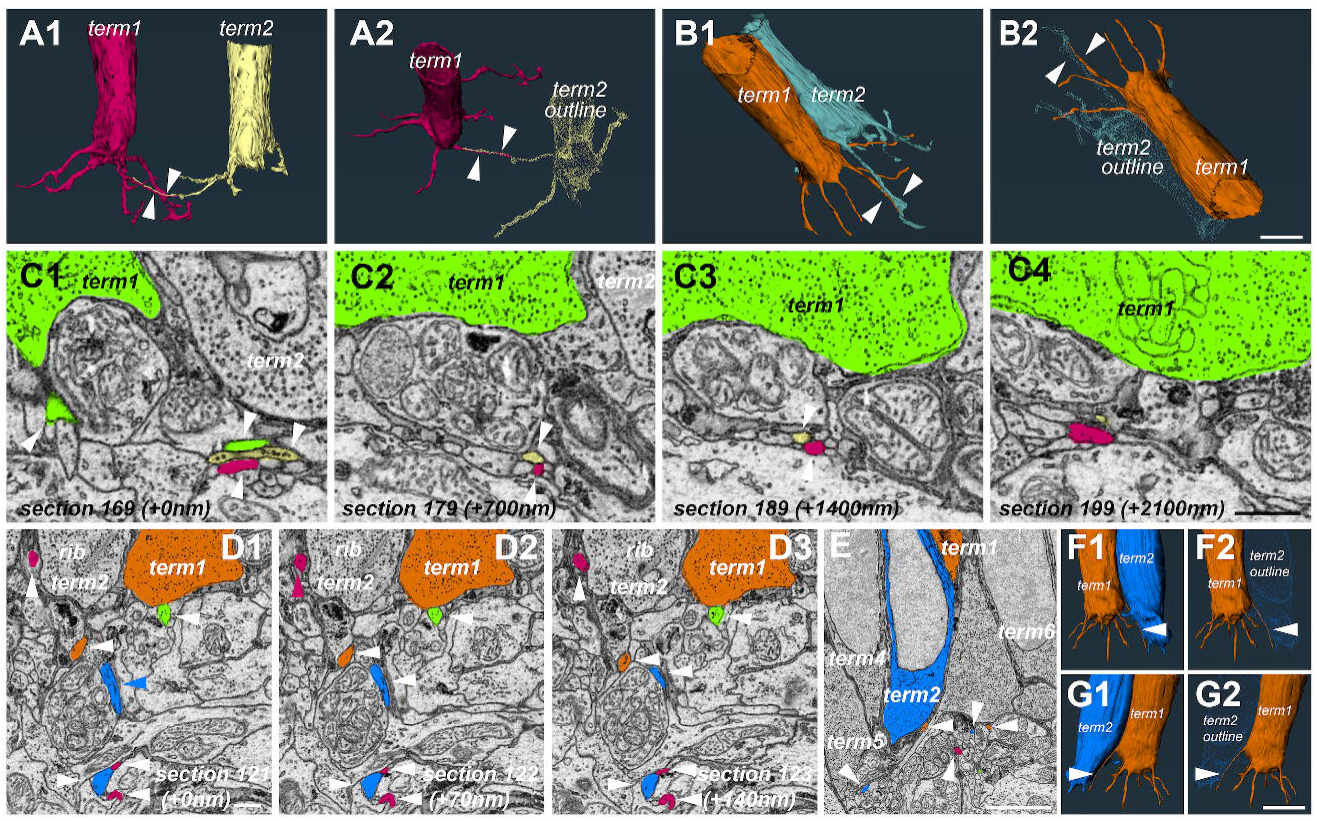
Telodendria association between non-adjacent and adjacent rods. A1-A2) 3D reconstructions of 2 terminals separated by ∼20µm from the P2R9 dataset showing their respective telodendria extending toward each other and running adjacent to each other (white arrowheads) for ∼10µm indicating a possible association via a non-chemical synapse connection. B1-B2) 3D reconstructions of 2 immediately adjacent rod terminals from the same dataset. Note the close proximity of both telodendria to each other (white arrowheads) and the lack of any other telodendria with such close association in these two terminals (scale bar = ∼5µm). C1-C4) EM images showing a progression of 700nm steps through the data (equal to 10 sections) and a demonstration of the close proximity (white arrows) of the telodendria from the photoreceptors in A1-A2. The telodendrion from the green terminal can also be seen in C1 running in parallel for a short distance. Possible synaptic vesicles in the yellow telodendrion can be seen in C1 as well. 30 sections later (i.e. ∼ 2100nm), the red and yellow telodendrion separate (scale bar = 1µm). D1-D3) A possible basal contact between the telodendrion of one terminal (green + white arrowhead) and the terminal of a non-adjacent rod (dark orange). Other white arrowheads show telodendria from neighboring photoreceptors crisscrossing the IPL; rib – synaptic ribbon, term – synaptic terminal (scale bar = 1µm). E) A telodendrion from a more distally positioned rod (term1, dark orange) extending past an adjacent rod (term2, blue) but remaining in close proximity to the soma suggesting possible contact. White arrowheads point to telodendria from the blue, dark orange and burgundy terminals (scale bar = 5µm). F1-F2, G1-G2) 3D reconstructions of the blue and dark orange terminals from different angles showing the proximity between the blue terminal and the dark orange terminal telodendrion (scale bar = 5µm).

The prevalence of synaptic vesicles along telodendria seems unusual, as they are assumed to mostly interconnect photoreceptors via gap junctions, not chemical synapses^55^. Unfortunately, we seem to lack the resolution in our current skate retina datasets to be able to confidently identify gap junctions between rods, or between the telodendria of different rods.

### Telodendria of different rods often run parallel to each other and come close to adjacent rod terminals to form putative basal contacts

Aside from the intricate, and likely connected, meshwork that rod telodendria formed, we occasionally encountered another unusual characteristic – namely, a telodendrion from one rod was in close apposition to the telodendrion of another rod, often over significant distances (Fig. 6). This could be observed in rods that are immediately next to each other (Fig. 6B1-B2), or rods that were separated by 20 or more micrometers from each other (Fig. 6A1-A2, 6C1-C4).

Occasionally, we could also identify putative synaptic vesicles in some of these processes (Fig. 6C1). As mentioned in the previous sections, our current data does not have enough resolution to definitively identify gap junctions, but we suggest that the close proximity of telodendria from adjacent and non-adjacent rods is not random, but rather suggestive of contacts. We also observed instances of close association resembling basal contact between the extending telodendrion of one rod and the terminal base of a non-adjacent rod, with a flattening of membranes between the telodendrion and the terminal, strongly suggesting association (Fig. 6D1-D3). Telodendria from more distally located rods also appear to be in close apposition directly to the terminal membrane of an adjacent rod for extended distances, suggesting contact from the adjacent rod (Fig. 6E, 6F1-F2, 6G1-G2). It is also possible that the telodendria of some rods invaginate into the terminals of adjacent rods, as has been observed in zebrafish^56^.

Although we have had several preliminary observations (data not shown) that suggest this might be happening for skate rods as well, we have not been able to confidently confirm that this is the case.

### Multiple post-synaptic processes invaginate into a single skate rod terminal forming structures that are unlike the typical tetrad observed in the terminals of rods from duplex retinas

Processes from vertebrate post-synaptic retinal neurons, namely bipolar cells and horizontal cells, tend to invaginate into the synaptic terminal of their target photoreceptor, be it rod or cone (for some of many examples see ^30, 46, 57–59^). This anatomy tends to be somewhat stereotypical and in mammalian rod terminals, often called spherules, four invaginating processes come from two horizontal and two bipolar cells, forming a so called “triad” or “tetrad” synapse^60–63^. These processes terminate at different invaginating depths under a single ribbon with the horizontal cell dendrites almost invariably closer to the ribbon and the bipolar cell dendrites more distant. However, variations and exception on this theme in mammalian retinas are beginning to be described^64^. On the other hand, cone terminals, often called pedicles, display a seemingly quite different anatomy, with multiple ribbons and multiple invaginating processes^47^. Upon closer examination, it becomes apparent that each synaptic ribbon bears a striking resemblance to a single rod spherule - i.e., there are two bipolar cell processes and two horizontal cell processes terminating under each individual ribbon. The whole cone pedicle, then, resembles a number of spherules brought together in a single terminal. In non-mammalian retinas, the individual anatomy of rod and cone terminals is similar, but there are some differences, depending on the species. For example, multiple ribbons seem to be present in rod terminals of amphibian rods^30^ and cone pedicles have a single large invagination^48^. Surprisingly, our very close inspection of the skate rod terminal revealed a different organization. Some of it we have already described in the previous sections (see section describing ribbons). What we noticed here, is the presence not only of multiple ribbons in each terminal, but also of multiple invaginating processes (Fig. 7B and 7E). Each rod terminal we examined did not have 4 invaginating processes, as would be expected from other work examining vertebrate rod terminals, but a much higher number, all clearly invaginating into that terminal. The partially reconstructed invaginating post-synaptic process of two neighboring rod terminals from the P2R9 volume are shown in Fig 7A1-A2 and Fig 7D1-D2. The individual post-synaptic processes for each terminal are shown individually in Fig. 7C1-C11 and Fig. 7F1-F10.

**Figure 7.**
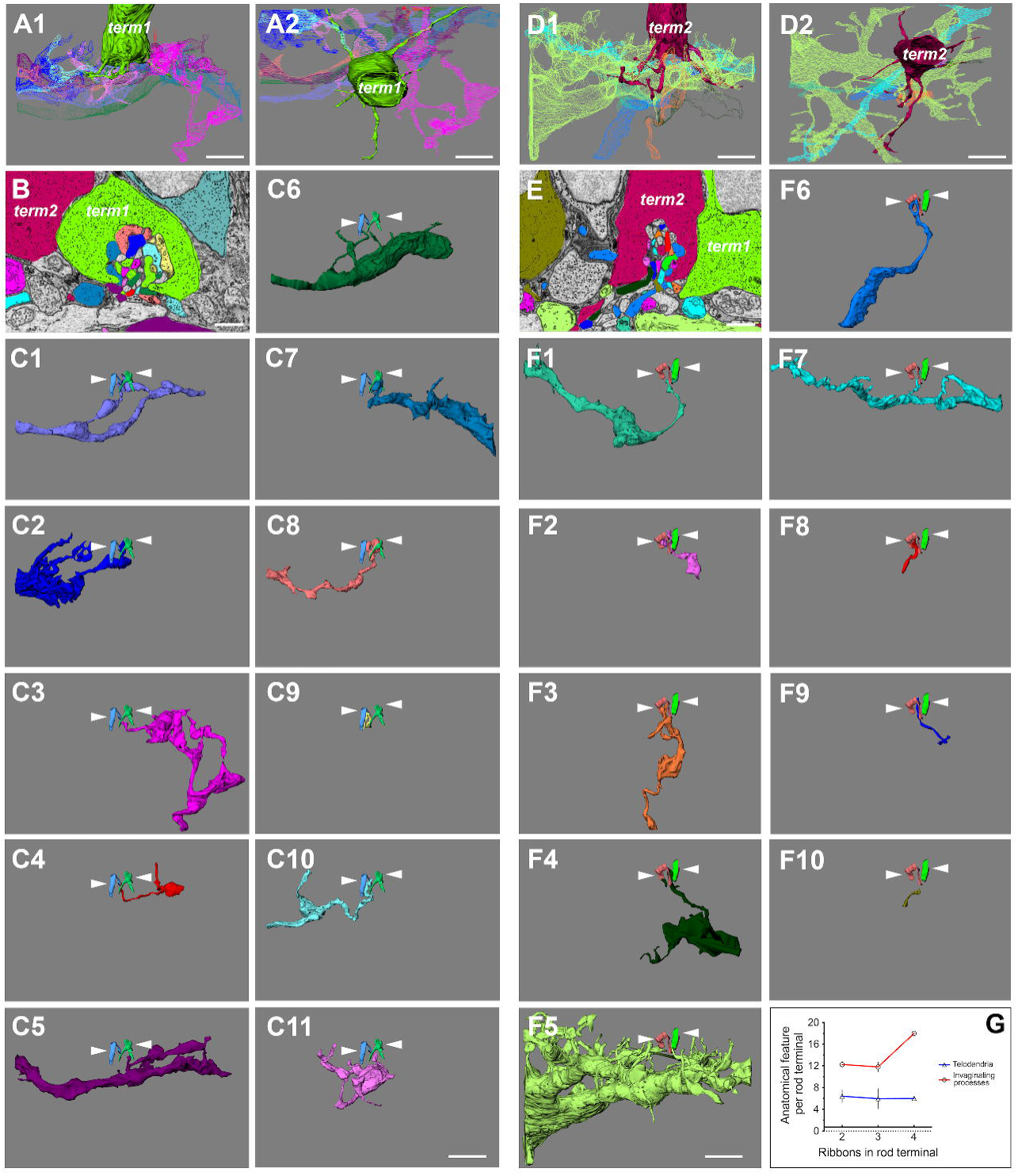
Multiple post-synaptic processes invaginate into a rod terminal. A1-A2) Side and top-down view of a partial rod terminal from the P2R9 dataset along with all of its reconstructed invaginating post-synaptic contacts (scale bar = 5µm). B) EM image and segmentation example of the same terminal (term1, green) and its invaginating processes (scale bar = 1µm). 1-11) 3D reconstructions of each invaginating process with the rod terminal removed for clarity. White arrowheads point to the 2 synaptic ribbons of the green terminal. At least 11 distinct processes appear to invaginate into this terminal (scale bar = 5µm). C1-C2) Side and top-down view of a partial rod terminal from the P2R9 dataset along with all of its reconstructed invaginating post-synaptic contacts (scale bar = 5µm). D) EM image and segmentation example of the same terminal (term2, burgundy), and its invaginating processes scale bar = 1µm). 1-10) 3D reconstructions of each invaginating process with the rod terminal removed for clarity. White arrowheads point to the 3 synaptic ribbons of the burgundy terminal. At least 10 distinct processes appear to invaginate into this terminal. B and D show that these two terminals are adjacent to each other but do not appear to share any of the reconstructed processes, with the exception of process 5 of term2 (scale bar = 5µm). E) A graph of the number of post-synaptic processes we have segmented from different terminals. There is no appreciable correlation between the number of ribbons and the number of invaginating processes for any given terminal (data for terminals with 4 ribbons is insufficient for any conclusions (n=1). Rod telodendria are also not significantly different between terminals with different number of ribbons and different number of invaginating processes.

Our quantifications show that there are at least ten, but as many as 18, distinct invaginating processes that we could reasonably assign to reconstructed post-synaptic partners. This is likely a conservative estimate, since we sometimes had to omit processes we thought were not sufficiently traced to establish them as anatomically distinct. Interestingly, there does not seem to be any correlation between the number of telodendria per rod terminal, and the number of invaginating processes (Fig. 7G). Due to the rare occurrence of terminals with 4 ribbons, we only have one such terminal with reconstructed invaginating processes, where there appear to be 18. This is insufficient data for us to make any definitive conclusions about a correlation between the number of ribbons and the number of invaginating processes for terminals with 4 ribbons and further investigation is necessary. However, terminals with 2 and 3 ribbons appear to have very similar number of invaginating processes (n=4, mean=12.25 for 2 ribbons; n=5, mean 11.80 for 3 ribbons; Fig. 7G). We currently do not have reliable data for terminals with 1 ribbon, as they are also rare and additional imaging is needed to capture all invaginating processes in such terminals.

## Discussion

In this study, we report a number of ultrastructural hallmarks of the functionally plastic rod photoreceptors of the skate retina. Using serial EM imaging, we show that skate rods display typical rod characteristics in their outer segments, but hybrid rod-cone characteristics in their inner segments and synaptic terminals. Thus, the skate rod almost appears to be separated into two distinct anatomical domains. Namely, rod outer segment architecture displays stacked membrane discs physically separated from the outer membrane, likely reflective of the physiology and ability of skate rods to function with great sensitivity at scotopic light levels^13^. On the other hand, their synaptic architecture appears to borrow elements of cone pedicle design, perhaps reflective of their ability to recover functionality under photopic light conditions^26^.

### The Little skate as a novel model in the study of the comparative neuroscience of vision

The physiology and molecular landscape of the duplex (i.e., mixed rod-cone) vertebrate retina have been worked out in great detail in the past ∼50 years, especially for mammals^65–68^. Transgenic approaches in animal models have also contributed greatly to our study of different elements of rod and cone circuitry, together, or in isolation^69, 70^. However, a large number of studies of the visual system have been performed on a fairly limited number of model organisms, like mouse, rat, rabbit, zebrafish, or salamander. This is likely because technical approaches have already been well established for these organisms, but it is worth noting that the almost exclusive study of the visual system in such model organisms has perhaps given us too narrow a focus and may have introduced bias in our understanding of broad and comparative principles of retinal design. The simplex, pure-rod skate retina provides us with an important new comparative model and an exciting avenue to study vertebrate rod circuitry within the context of a functional and evolutionarily optimized system. Furthermore, the ability of this simplex retina to function under both scotopic and photopic ranges of illumination gives a unique opportunity to examine how complete our understanding of retinal design and physiology is. Indeed, only in the last several years, a long-held assumption that rods saturate and are mostly inactive under photopic conditions in a duplex retina has been challenged^71, 72^. Thus, such novel comparative models should not be underestimated, as they have the potential to add exciting new avenues in vision restoration efforts and to aid our overall understanding of the vertebrate visual system.

### Similarities and differences between rods from mixed rod-cone duplex retinas and rods from the simplex of all-rod skate retina

#### Outer segments, inner segments and synaptic terminals

Our detailed examination of the outer segments of skate rods using a SB-3DEM approach has easily confirmed the typical architecture of stacked membrane disks observed previously by Szamier and Ripps^18^, and typically expected of vertebrate rods from mammalian and non-mammalian retinas, alike^73^. This architecture is different from vertebrate cones, which have an outer segment membrane and stacked membrane disks that are continuous with each other^74^. These outer segment structural hallmarks were long thought to be important mediators of function in both rods and cones, but an elegant recent study in lamprey retina by Morshedian and Fain definitively showed that the single-photon sensitivity of rods is not intimately connected to the separation of stacked membranes in the outer segments of rods, since lamprey rods have cone-like outer segments^75^. Skate rods appear to have sensitivity approaching single photon detection^13^ and despite their functional plasticity and ability to light-adapt to photopic levels of illumination, they display a strikingly stereotypical outer segment morphology, as we have confirmed here. Yet, strikingly, they still retain an ability to speed up their kinetics, lower their sensitivity, and expand their functional capabilities to cone levels of illumination. We propose that some of that remarkable capability has structural underpinnings manifested in the inner segment and synaptic terminals of skate rods. We find, for example, that skate rods have inner segments rich in mitochondria displaying mixed rod-cone hallmarks (data not shown here; in preparation for a separate report). In the results presented here, we show that skate rod terminals have a varying number of ribbons (between 1 and 4, and possibly more) and that there is a skewed distribution toward 2-3 ribbons/terminal, among the rod terminals we examined. We cannot say as yet what functional consequence a different number of synaptic ribbons per terminal may have, but additional data (not presented here) points to a non-selective connection of putative bipolar and horizontal cells to rods with different number of ribbons. In fact, it appears that neurons post-synaptic to rods might make connections to rods based on what is available in their dendritic field, rather than selectivity based on a specific rod attribute. As mentioned before, previous literature indicates that other vertebrate species, like teleost fish^59, 76^ and mammals^35, 77^, have a single synaptic ribbon per rod, although exceptions for mammals have been described^78^. Amphibians, on the other hand, tend to have multiple ribbons in their rods^30, 79, 80^. However, systematic studies of rod synaptic ribbons in different vertebrate retinas are largely lacking and a description of these organelles is often in a functional context, or in the context of describing structural motifs of connectivity. To our knowledge, a systematic quantification of ribbon distribution across rod photoreceptors in a simplex retina, as we have performed here, is completely lacking.

#### Telodendria with synaptic vesicles and gap junctions

The multiple telodendria extending from each skate rod terminal were another intriguing finding from our study. Telodendria appear to be mostly associated with mammalian and non-mammalian cones^49, 53^, where they are largely believed to be sites of cone-to-cone gap-junctional contacts^55^. Mammalian rods seem to lack telodendria^54^, but at least some non-mammalian rods have extensive telodendria networks^56^. Our results also strongly suggest that conventional chemical synapses along rod telodendria are present (see Fig. 5). This type of architecture might be in place of, or in addition to, gap junctional contacts along those same telodendria. At present, we do not have sufficient resolution in our data to exclude or confirm the presence of gap junctions along these processes, or between the terminal endings of adjacent rods. Nevertheless, we suggest that the role of rod telodendria in skate retina is to indeed mediate some form of receptor coupling and therefore possibly improve sensitivity^81^, either through chemical or electrical synapses, or both.

#### Invaginating contacts

Yet another surprising finding in this study has been the number of individual post-synaptic processes we were able to identify in the single invagination of skate rods (see Fig. 7). The high number of invaginating processes (mean of 12.85 processes per rod terminal) is not typical of rods, yet the single invagination into the terminal is. For example, mammalian rods typically have a single invagination with 4 dendritic processes (evenly split between horizontal and bipolar cells) terminating proximally and distally to the synaptic ribbon, respectively^43^. On the other hand, mammalian cone pedicles tend to have multiple ribbons (exact number depends on the cone type) and multiple invaginations, each with ∼4 invaginating processes^46^. This arrangement is somewhat less organized in non-mammals, but generally, there are fewer invaginating process in rods^30, 45^ and more in cones^82–84^. To our knowledge, the ultrastructural characteristics of synaptic terminal arrangement of photoreceptors in a simplex retina has never been described.

### Skate rods and downstream circuitry exhibit hybrid rod-cone architecture

In closing, our observations in this study are consistent with assigning a “hybrid” rod-cone characteristic to skate rods. This conclusion is supported by the presence of a number of uncharacteristic structural features, including multiple ribbons centered over a single rod synaptic invagination, long extending telodendria that form intricate networks, and multiple invaginating contacts into each rod terminal. We suggest that all of these features are likely part of the structural underpinnings of functionally plasticity in skate retinal circuitry.

## Acknowledgements

This work was supported by National Institutes of Health grants SC2-GM144198 (to I.A.A.), MBRS-RISE R25-GM059298 (to L.M.H and P.E.P.), MS/PhD Bridges R25-GM048972 (to Y.Y.), California State University Program for Education and Research in Biotechnology New Investigator Grant (to I.A.A.), and by the Genentech Foundation Scholarship (to J.G.F. and B.R.). We would like to thank Grahame Kidd and Emily Benson for help with EM data acquisition and helpful analysis suggestion, Richard Chappell for logistical support at the Marine Biological Laboratory, David Berson, David Copenhagen and Felice Dunn for helpful discussions and technical support, Altan Od Baatar for support with pilot segmentation, Karl Murphy for expertise and technical support with animal husbandry, and Robyn Crook for help with statistical analysis, animal husbandry, and comparative neuroscience discussions.

